# Unlocking Protein Evolution Insights: Efficient and Interpretable Mutational Effect Predictions with GEMME

**DOI:** 10.1101/2024.08.15.608048

**Authors:** A. Carbone, E. Laine, G. Lombardi

## Abstract

The Global Epistatic Model for predicting Mutational Effects (GEMME) is a computational method that reconstructs protein mutational landscapes using sequence data alone. In this article, we delve into the broader biological questions that GEMME can address beyond mere landscape reconstruction. We provide several examples to guide users in maximizing the utility of GEMME. Additionally, we discuss recent advancements that enhance GEMME’s predictive accuracy and extend its capabilities by integrating sequence data with structural information and allele frequency data, when available. These innovations enable GEMME to answer new biological questions and offer deeper insights into protein evolution and function.

## 1 Introduction

The systematic and accurate assessment of the impact of mutations on protein functioning is of critical importance in biology, bioengineering, and medicine. The Global Epistatic Model for predicting Mutational Effects (GEMME) is a computational tool that reconstructs these landscapes using only protein sequence data across species, significantly contributing to recent advancements in this field [1].

GEMME is an unsupervised and rapid method that predicts the effects of mutations by explicitly modeling the evolutionary history of natural sequences. It employs a few biologically meaningful and interpretable parameters and globally considers epistasis by evaluating the whole sequence context when assessing a mutation’s impact. Given an input alignment, GEMME can generate a protein’s full mutational landscape by assessing the impact of all possible single amino acid substitutions in just a few minutes. It performs on par with high-capacity deep learning-based architectures [2]. It is applicable to both single-site mutations and combinations of mutations. It is freely available for the community through a stand-alone package and a web server at http://www.lcqb.upmc.fr/GEMME.

GEMME proved useful for discovering functionally important residues in proteins and distinguishing them from residues essential to protein thermodynamic stability [3, 4, 5]. It helped to decipher the molecular mechanisms underlying various diseases [6, 7] and to estimate mutation-induced binding affinity changes in protein-protein interactions [8]. Moreover, using GEMME scores as surrogates for experimental measurements allowed for boosting protein language model (pLM) accuracy and speed [9] with the resulting predictor, VespaG, combining the best of both worlds, the robustness and transparency of GEMME with pLM universal protein representation space.

This contribution explores broader biological questions that GEMME can address beyond landscape reconstruction by providing four examples to help users maximize GEMME’s utility. They are presented with a fine-tuned version of GEMME, named iGEMME, with optimized parameters enhancing its performance [10].

We also highlight recent advancements, ESCOTT and PRESCOTT [10], which improve GEMME’s predictive accuracy and extend its capabilities. ESCOTT integrates sequence data with structural information, while PRESCOTT incorporates both structural information and allele frequency data when available. These innovations enable GEMME to answer new biological questions and provide deeper insights into protein evolution and function.<H

### 1.1 Biological questions GEMME can help to address

Multiple questions arise directly from mutational landscape reconstructions:

Do mutational landscapes provide insights into which regions along the protein sequence are more sensitive to mutations than others?

When structural data is available, do mutational landscapes indicate which three-dimensional regions of the protein are more sensitive to mutations?

Can GEMME be used to discover and annotate sites in protein sequences with distinguished functional and structural roles?

Can GEMME identify key regions in intrinsically disordered proteins that might suggest interaction sites?

More broadly, what are the mechanisms underlying disease mutations at a proteome-wide scale?

Is there a correlation between regions predicted to be sensitive by GEMME and regions where tools reconstructing the structure, like AlphaFold [11], ESMFold [12] and RosettaFold [13], provide confident predictions?

Are regions that AlphaFold, ESMFold, RosettaFold predict with uncertainty discovered to be sensitive to mutations by GEMME? In other words, can GEMME provide complementary information about a protein structure that structure predictors do not reveal?

How can we visualize sensitive regions within a protein sequence? and on a protein structure, whenever available?

Can GEMME model go beyond one mutation? Is GEMME model useful to predict the effect of two or more mutations?

Based on the analysis of four distinct proteins, we will guide the user in utilizing GEMME to answer to these questions: the thioredoxin TRX is a widespread protein family with a highly conserved structure across the tree of life [14, 15, 16]; the homeobox protein ARX is a disordered protein with no structural prediction available from AlphaFold [17, 18]; the chromodomain of the heterochromatine protein 1 HP1, a major component of heterochromatin, directly binds to H3K9me [19, 20, 21]; the nuclease NucB is an enzyme used to explore generated sequences for designing optimized proteins in various environments [22, 23].

## 2 Methods

### 2.1 GEMME/iGEMME reconstructs mutational landscapes of proteins: input and output

GEMME/iGEMME uses a protein sequence as a reference to provide a mutational landscape for that specific sequence. The computations are based on a multiple sequence alignment (MSA; Figure 1B) with the reference sequence on top. No gaps are allowed within the reference sequence (columns in the MSA with gaps in the reference sequence should be deleted), such that the MSA is of the same length as the query sequence, namely *L*. By default, GEMME/iGEMME generates an *L* × 20 matrix of scores, with 20 the size of the amino acid alphabet, representing the full single-site mutational landscape of the reference. It can also provide predicted scores for a user-defined list of single or multiple mutations.

**Figure 1:**
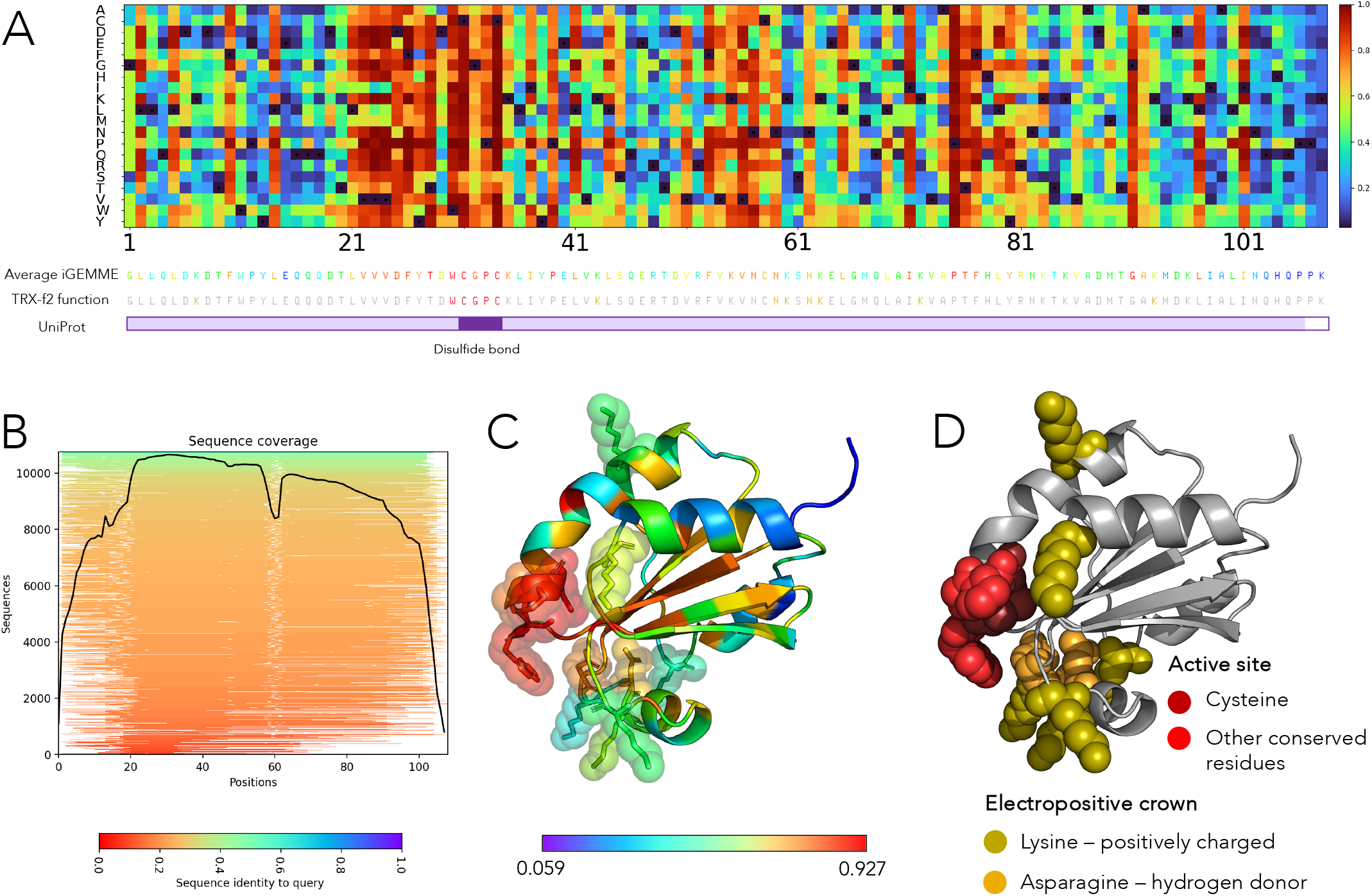
iGEMME analysis of the thioredoxin, type f, TRX-f2. **A.** iGEMME matrix of TRX-f2. iGEMME scores are ranksorted from high effect (red) to no effect (blue) based on the color bar on the right. Residues associated to the 108 TRX-f2 sequence positions are colored with the average score of the columns in the iGEMME matrix. Residues known to be functionally important for the TRX-f2 protein are reported with colors as in the legend in C, and all others are in grey. Uniprot (ID A0A2K3DSC9) domain annotation with the TRX domain highlighted in light purple and the disulfide bond motif in dark purple. **B**. ColabFold profile of sequence coverage (black line) for the MSA used by iGEMME to predict TRX mutational effects. The sequence identity to the query is described by the colors. **C**. TRX-f2 structure (PDB 6i1c) reports average iGEMME scores. Same colors scale as in A. **D**. TRX-f2 structure reports residues known to play an important biological role for TRX-f2: the extended disulfide bond motif (WCGPC) in red tones, positively charged crown surrounding the active site, comprised of lysines and asparagines, in orange tones. All other residues are left grey. The two cysteines in the WCGPC motif, colored in dark red, are positioned on the back of the red motif on the left hand side of the structure and are barely visible.

Figure 1A shows the iGEMME mutational landscape matrix for the thioredoxin protein TRX of type f from *Chlamidomonas reinhardtii*. The iGEMME scores, corresponding to single mutations from the reference sequence, are represented on a color scale ranging from red (indicating highly sensitive mutations) to blue (indicating no mutational effect).

Several positions in the matrix exhibit red tones for most or all mutations at that position, indicating highly sensitive sites. These sites are often grouped in consecutive regions and typically correspond to structured areas with potential functional roles. They could also be located on small loops or be indicators of the formation of transient structures. In contrast, positions displaying blue tones indicate areas with no mutational effect, likely in parts of the protein that remain unstructured throughout its lifetime, but not only, as illustrated by the blue helix in Figure 1C.

GEMME/iGEMME scores averaged per column reflect residues’ sensitivity to mutations. Hence, a useful way to analyze GEMME/iGEMME results and capture positional relevance in the global context is to map these average values on an experimental or predicted three-dimensional (3D) structure for the reference protein sequence. This approach allows for identifying 3D regions sensitive to mutations. See Figure 1A for an example of average scores associated with residues in the sequence, and Figure 1C for the plot of these averages on the structure. Sensitive residues are highlighted in red in both panels.

### 2.2 Recognition and discovery of positions and regions highly sensitive to mutations

Analyzing the average sensitivity of positions in a sequence or structure with GEMME/iGEMME provides biological insights into 2D/3D regions that are more mutation-sensitive than others. For instance, in Figure 1, the known disulfite bond motif at positions 102-105 (CGPC) is identified by iGEMME as highly important (red) for all thioredoxins. Specifically, iGEMME identifies the extended motif WCGPCK as highly significant (red residues in Figure 1D), supporting existing observations that additional residues are involved in an extended active-site three-dimensional motif controlling the reactivity of the thioredoxin fold [24]. Key residues are located on the beta sheets in front of the motif, which iGEMME identifies as highly sensitive regions.

For the specific TRX-f2 protein, it has been observed [25] that the disulfite bond motif is surrounded by a crown of lysines (Lys7, Lys43, Lys60, Lys63, Lys72, Lys93; olive green residues in Figure 1D) and asparagines (Asn59, Asn62; orange residues in Figure 1D). Most of these residues are identified as important (score > 0.4) by iGEMME. Compare Figure 1D with Figure 1C, where scores are represented by residue colors, ranging from red to green.

GEMME/iGEMME can also help to discover new regions of the protein that have not yet been identified with biological significance. Several regions in the TRX-f2 sequence are highlighted by iGEMME as important (red regions in Figure 1C) but these regions have not yet been mentioned in the literature. In conclusion, by inspecting the global matrix of mutational effects, iGEMME can help discover important functional sites that have not been previously annotated in databases like UniProt [26] and InterPro [27], and guide the design of mutational experiments to explore these sites further.

### 2.3 Finding functional motifs in intrinsically disordered regions or regions with few homologs

Intrinsically disordered proteins or disordered regions within proteins are expected to interact with other molecules, including structured proteins [28, 29]. During complex formation some transient structures may form. The precise location of these transient motifs, which help in binding with partners, remains an open question. GEMME/iGEMME can likely contribute to this area of study. These ephemeral regions are thought to be encoded in sequences that meet specific physico-chemical and geometrical conditions, which should be detectable by evolutionary sequence constraints.

As illustrated in Figure 1, relevant sensitive sites can be highlighted using iGEMME matrix, and new suggested motifs can be further annotated to understand protein function. This is particularly important when no structural model is available for the protein, as shown for the protein ARX, which is associated with mental retardation and epilepsy [30], in Figure 2. Annotations from Uniprot indicate that ARX exhibits a high degree of disorder primarily concentrated in the initial segment of the protein (1-255 in Figure 2A), while the folded portion encompasses the homeobox domain (328-390) and the OAR region (530-543). iGEMME identifies distinct signals of sensitivity within the disordered regions, indicating potential structure formation. Specifically, these signals correspond to residues 1-21, 50-81, 157-181 in the initial segment, and also between the homeobox and OAR domains (391-529). These regions are highly disordered in the AlphaFold model (Figure 2C). The prediction provided by ESMFold shows a more structured pattern (Figure 2D) than AlphaFold, but with comparable confidence for both predictors (Figure 2E). Thus, GEMME/iGEMME can serve as a valuable complementary tool to state-of-the-art protein structure predictors.

**Figure 2:**
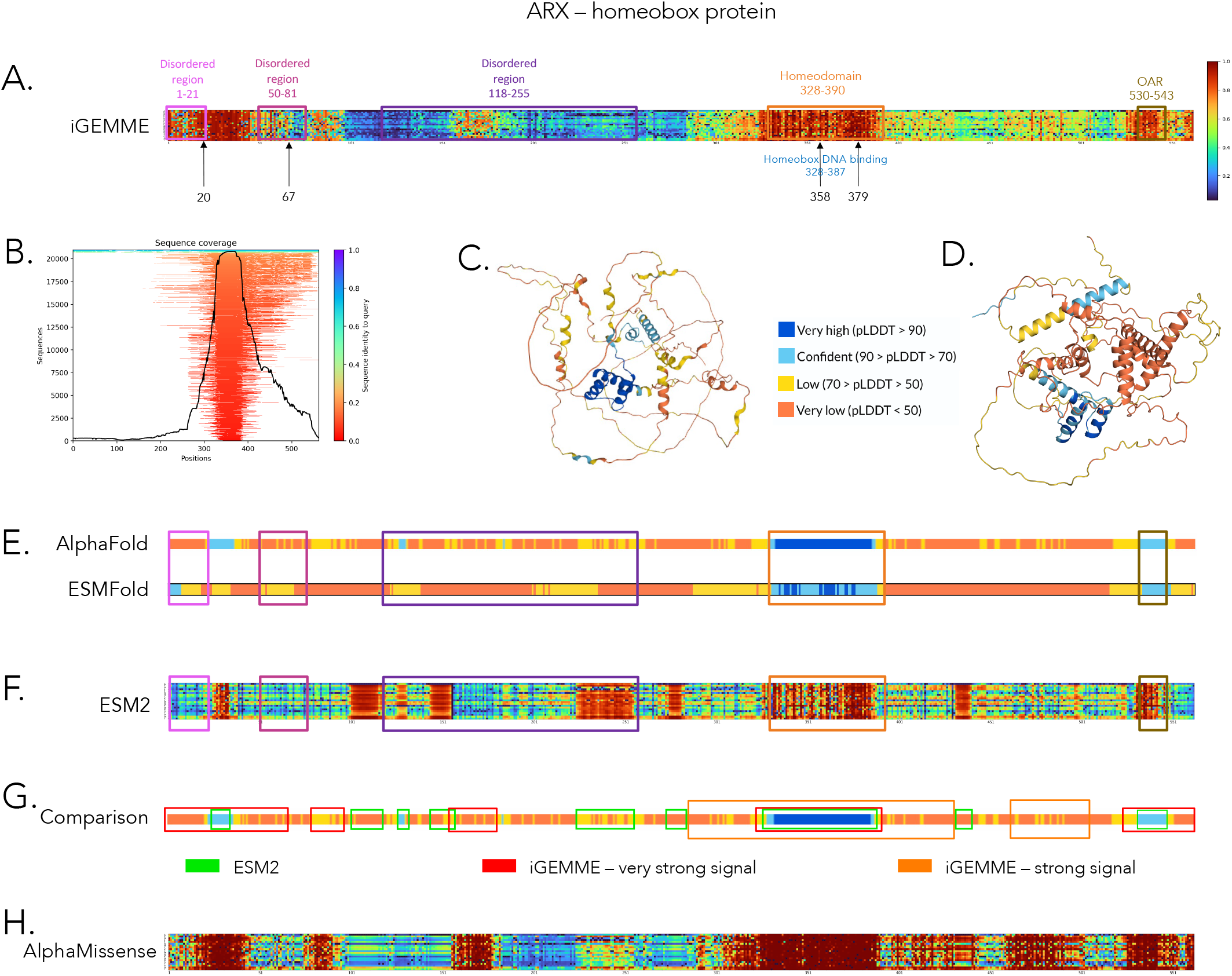
The human homeobox protein ARX analysed with iGEMME and other methods. **A.** iGEMME matrix is annotated with domain information coming from UniProt (www.uniprot.org/uniprotkb/Q96QS3/entry). Residues 20 and 67 are annotated as phosphoserines. Positions 358 and 379 in the homeobox domain are known to abolish sequence-specific DNA-binding, to reduce transcriptional repression of LMO1 and SHOX2, and abolish transcriptional activation of the enzyme KDM5C. iGEMME scores are ranksorted. **B**. ColabFold profile of sequence coverage (black line) for the MSA used by iGEMME to predict ARX mutational effects. The sequence identity to the query is described by the colors. **C**. AlphaFold2 3D model, downloaded from UniProt, see B. **D**. ESMFold 3D model, computed locally. **E**. Positional confidence of the AlphaFold2 and ESMFold predicted structures. For each residue along the sequence, confidence is measured by pLDDT scores grouped in four main classes: high (blue), medium (light blue), low (yellow) and very low (orange). AlphaFold prediction is retrieved from UniProt (www.uniprot.org/uniprotkb/Q96QS3/entry). ESMFold prediction has been computed locally. Domain annotation is reported from A. **F**. ESM2 zero-shot predictions of ARX mutational effects. ESM2 scores are ranksorted. **G**. The AlphaFold2 confidence bar in B is aligned with regions in iGEMME (red boxes for very high scores and orange boxes for high scores - see red and orange colors in A) and in ESM2 (green boxes; see red colors in E). **H**. AlphaMissense prediction of ARX mutational effects. Compare with panel A.

A recent study used ARX to illustrate the ability of the protein language model ESM1b to detect signals independently of MSAs, allowing it to transfer information from well-covered regions in proteins to poorly covered ones [31]. However, we found that GEMME/iGEMME can recover similar signals directly and solely from an input MSA, even if the latter does not uniformly cover the sequence and includes regions with very reduced coverage or highly divergent sequences (Figure 2B). This finding is further supported by the comparison of iGEMME results with the zero-shot scores computed with the pLM ESM2 (650M parameters) [12], which are derived from log-odds ratios of predicted amino acids probabilities for each position (Figure 2G). Both iGEMME and ESM2 managed to pick up signals in disordered regions or regions with few homologs and the identified sensitive regions overlap to some extent.

Hence, this example illustrate the ability of GEMME/iGEMME to detect signals even in regions where the available sequence data is limited (Figure 2B). These results are consistent with our previous studies emphasising the robustness of GEMME with respect to low MSA variability [32, 10, 1] and its high performance on viral proteins [2, 1]. This is a significant finding because, given the speed of GEMME/iGEMME and the rapid generation of MSAs with ColabFold [33], the advantage of using protein language models to predict mutational effects becomes less clear.

We also specifically want to highlight that GEMME’s performance can be directly compared with more data-intensive tools like AlphaMissense [34], which relies on the AlphaFold model and incorporates extensive additional information including structural data and allele frequencies. This comparison is evident when examining Figure 2G (red and orange regions) alongside Figure 2H (red regions), demonstrating that iGEMME, which uses only an MSA, can achieve similar results despite the more limited data input.

### 2.4 Identification of residues essential for protein functioning and not folding

Contrasting GEMME/iGEMME evolutionary-based analysis with complementary information about thermodynamic stability can reveal functionally important residues not essential for folding. We illustrate this concept with the chromodomain of HP1 (Figure 3) for which folding stability measurements of 90% of all possible single mutants are available [3]. Computed from an input MSA containing over 10 000 diverse sequences (Figure 3A), GEMME mutational landscape (Figure 3B) uncovered four residues, namely F2, Y26, D30, and E34, whose evolutionary constraints suggest a functional role while they are unimportant for thermodynamic stability (Figure 3C, orange dots). These residues are essential for the binding of methylated histone H3 (Figure 3D, left panel), either by establishing a direct contact or by forming a cage into which the methylammonium group inserts [35]. In addition, GEMME identified six residues highly sensitive to mutations, namely I7, Y19, V21, W23, W33, F50, that are also crucial for the domain’s thermodynamic stability (Figure 3C, red dots). These residues are all hydrophobic and buried or part of well-defined secondary structures (Figure 3D, right panel). Overall, GEMME evolutionary scores are consistent with physical constraints, and none of the residues predicted as tolerant to mutations are structurally critical (Figure 3C). A more systematic comparison of GEMME scores and mutation-induced folding stability changes can be found in [3].

**Figure 3:**
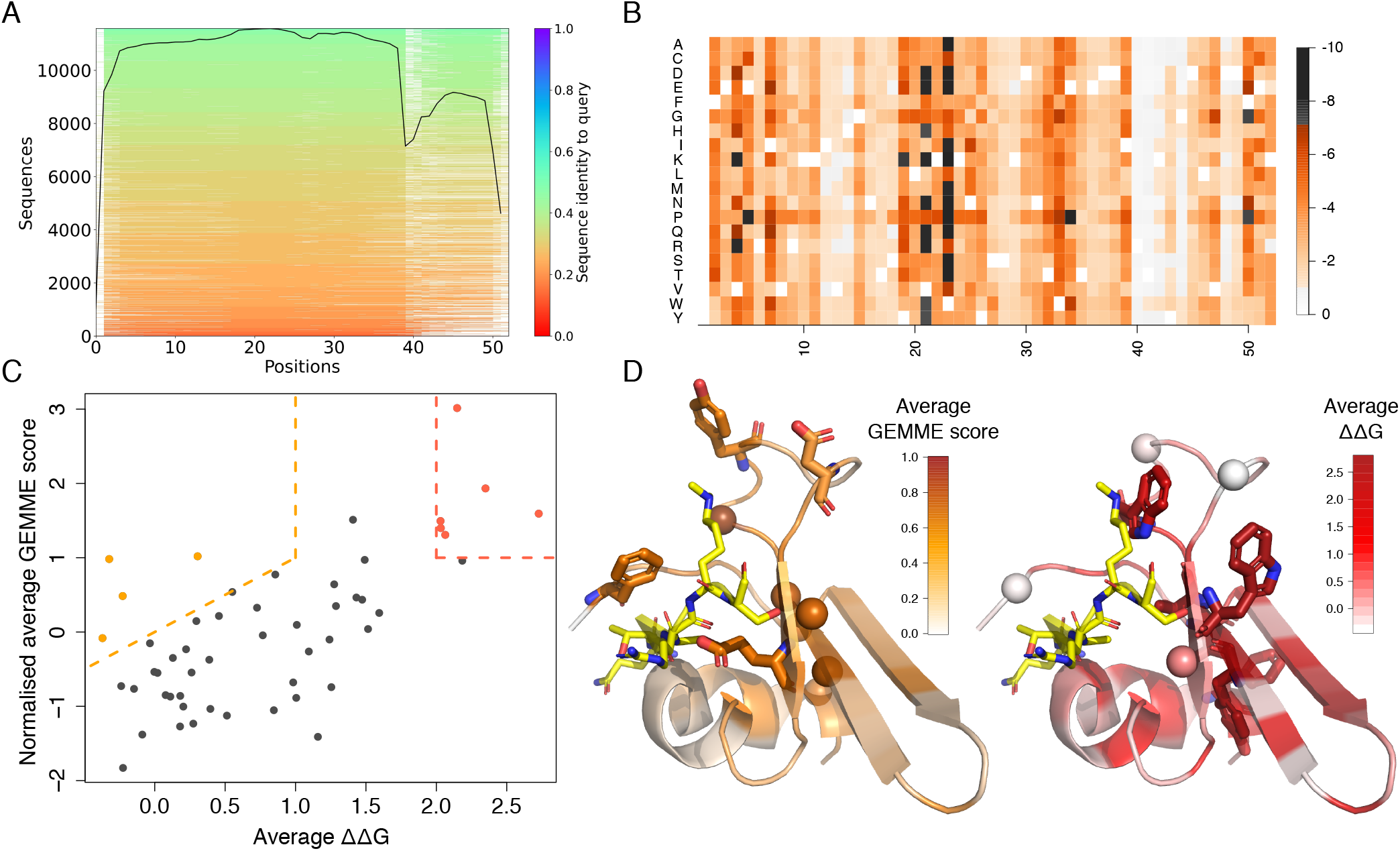
Comparison between evolutionary constraints and thermodynamic structural stability in the chromodomain of HP1. **A.** Alignment generated with ColabFold and given as input to GEMME. The sequence of the PDB entry 2M2L (residues 12 to 63) is used as query. **B**. Single-site mutational landscape computed by GEMME. **C**. Normalised average GEMME scores in function of average experimental ΔΔ*G* values taken from [3]. Each dot represents a residue/position. Normalisation consists in subtracting the mean and dividing by the standard deviation. Residues above the orange dotted line are sensitive to mutations according to GEMME/iGEMME but unimportant for stability; we refer to them as “functional” sites. Residues enclosed by the red dotted lines are sensitive to mutations from both evolutionary and structural perspectives. **D**. Mapping of average GEMME scores (left, in orange tones) and average ΔΔ*G* (right, in red tones) on the 3D structure 2M2L. The ligand, positioned by superimposing the structure 1KNA, is shown in yellow. On the left panel, functional residues are in sticks, and structurally critical residues detected as sensitive by GEMME are in spheres. Reciprocally, on the right panel.

### 2.5 Prediction of multiple mutations

GEMME/iGEMME is designed to predict multiple mutations by independently evaluating each mutation. While this model does not fully capture the interdependence of residue positions within a sequence, iGEMME’s performance remains competitive with many other zero-shot predictors, exclusively based on sequences. To demonstrate this, we analyzed one of the most extensive genotype-phenotype enzyme activity landscape to date. This dataset comprises 55 759 diverse variants of the nuclease enzyme NucB (github.com/google-deepmind/nuclease_design) [36]. The dataset has been used to optimize NucB for functioning in novel chemical environments, a key objective in synthetic biology. The goal is to show that models trained solely on evolutionary data, without any experimental data, can design functional variants at a significantly higher rate than traditional approaches to initial library generation.

The dataset comprises three rounds of mutational campaigns (G2, G3, G4), in which mutations where selected through a variety of techniques including direct evolution (DE), hit recombination (HR) and machine-learning models fit on evolutionary data and mutational data from previous generations (ML). G2, G3 and G4 campaigns were all built upon the same initial library (G1), obtained by applying error-prone PCR (epPCR) to a single-site saturation mutagenesis library of the wild-type sequence from *B. licheniformis*. Among more than 211K initial mutations generated by epPCR, employed for the DE campaign, only 9.4K where included in G1, corresponding to those for which catalytic activity was measured (Figure 4A). Based on their activity, mutated sequences where divided in 4 classes, namely *non-functional, activity >* 0, *activity > WT* and *activity > A73R*, the latter being the top-performing variant from the initial epPCR library, with 8-fold higher activity than the WT. iGEMME prediction of 2088 single-point mutations in this dataset is reported in Figure 4B.

**Figure 4:**
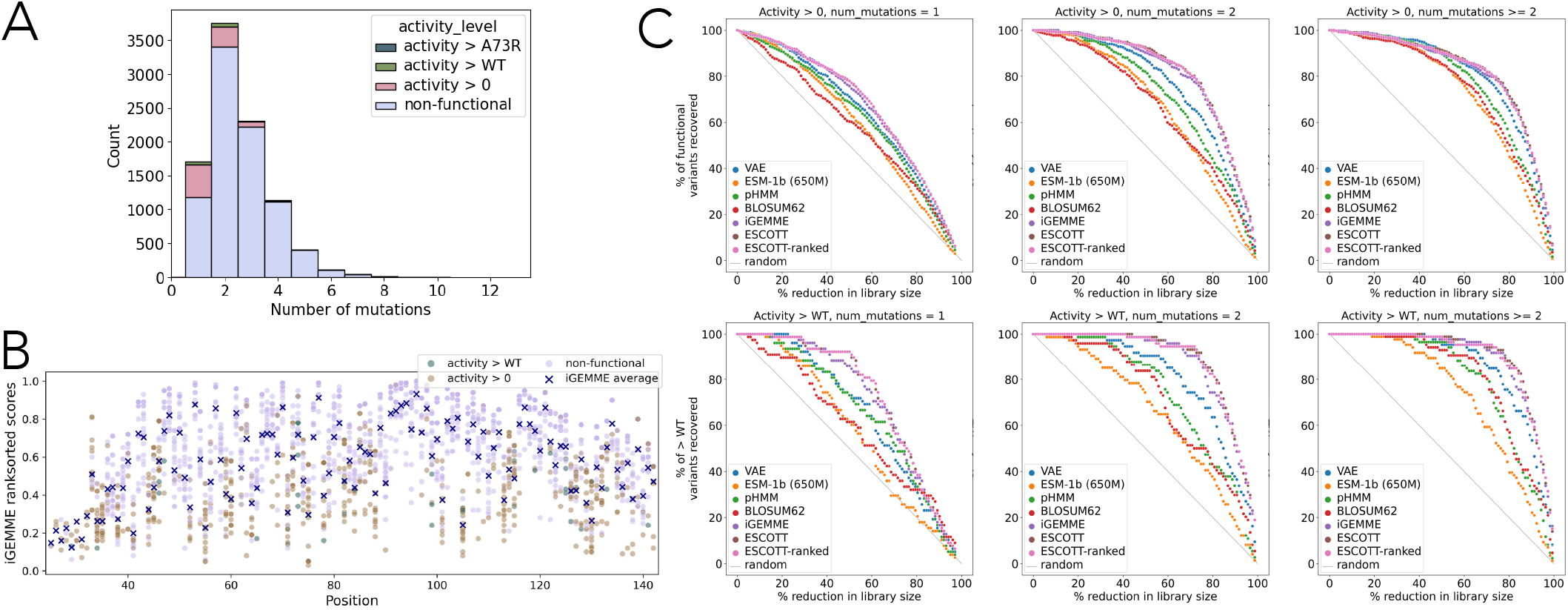
Comparative zero-shot analysis of the nuclease activity, NucB, with different tools predicting multiple mutations. **A.** Activity level distribution for the dataset of approximately 9 400 variants in G1 [36] colored by estimated activity levels. G1 includes a single-site saturation mutagenesis library as well as variants generated by error prone PCR. On average, G1 variants have 2.5 mutations, while functional variants have, on average, 1.6 mutations (from Figure 5f or Supplementary Figure 28 in [36]). **B**. iGEMME ranked sorted analysis of the 2088 single-point mutations present in the dataset. This analysis includes the approximately 1700 single-point mutations in G1 described in A. The prediction concerns three phenotypic classes of variants: with activity *> WT* (green), with activity *>* 0 (brown), non functional (light purple). For each position in the sequence, iGEMME averages are indicated by a cross. Low iGEMME scores indicate no mutational effect, hence functional activity, while high iGEMME scores indicate no functional activity. **C**. VAE, ESM-1b, pHMM, BLOSUM62, iGEMME, ESCOTT (raw scores) and ESCOTT (ranked) are compared on two tasks, for sequences with 1, 2 or more mutations: the percentage of functional variants recovered (top); the percentage of sequences that after mutation improve their activity with respect to the WT (bottom). The dataset used for this analysis is a subset of G1, previously used and described in [36]. See Supplementary Figure 6 in [36].

Figure 4C illustrates the superior performance of iGEMME compared to VAE [36], ESM-1b [37], pHMM [38], BLOSUM62 [39]. This comparison is based on two different evaluations: the percentage of functional variants recovered (Figure 4C, top) and the percentage of sequences that after mutation improve their activity with respect to the WT (Figure 4C, bottom). Notably, ESCOTT, which incorporates structural information into iGEMME, further enhances performance [10]. The analysis was conducted on a subset of the G1 variants: 874 have activity higher than 0, and 103 have an activity higher than WT (Figure 4A).

## 3 Notes: good practice and limitations

### 3.1 Generation of the input alignments

While GEMME initial design used two sets of aligned sequences as input [1], we advise users to give only one input MSA to the tool. This simplification improves interpretability without compromising on predictive accuracy. Moreover, as shown in [32] on a comprehensive benchmark, GEMME achieves similarly competitive performance with MSAs generated either by the highly efficient MMseqs2-based strategy implemented in ColabFold [33] or by highly sensitive profile Hidden Markov Model (HMM) workflows [40]. Users can generate an input MSA with the profile HMM-based method JackHMMer [40] against UniRef100 [41] via GEMME webserver, available at http://www.lcqb.upmc.fr/GEMME. The webserver also facilitates the import of alignments pre-computed by ColabFold, simplifying accessibility to both strategies.

The iGEMME results reported here were produced starting from MSAs computed by ColabFold, based on the uniref30 2103 and colabfold envdb 202108 databases of sequences. iGEMME was used through the webserver at http://prescott.lcqb.upmc.fr [10] without giving any structural model as input.

In the general case, we recommend users to run GEMME/iGEMME on MSAs obtained from Colab-fold with the most recent versions of the Uniref30 and environmental databases, particularly if they wish to perform high-throughput predictions. When the number of sequences retrieved is lower than 200, we recommend to remove ColabFold default filter [32].

### 3.2 Optimisation of GEMME parameters

The default parameters in iGEMME have been optimized compared to the original version of GEMME. The modified parameters concern the calculation of evolutionary conservation values by the JET algorithm [42]. Namely, the maximum number of sequences processed has been increased from 20,000 to 40,000, and the parameter limiting the number of sequences due to computational load (CPU load or max load parameter) has been raised from 500,000 to 8,000,000. Additionally, the maximum heap size has been expanded from 1,024MB to 8,192MB. These adjustments enable the handling of large proteins with MSA files containing thousands of sequences [10]. While users can tune these parameters based on their specific systems, the recommended values are generally suitable for most cases [10]. In addition, JET includes a Gibbs sampling-like procedure to subsample the input sequences. This stochasticity can generate different results from one run to another. To mitigate this effect, JET offers the possibility to run its algorithm several times (iJET) and retain the maximum conservation values over all runs. Our experiments with the tool indicate that setting the number of iterations to 7 leads to stable results. We thus recommend this value to users.

### 3.3 What might go wrong on GEMME/iGEMME

Some common issues include having a very long reference sequence (> 3 500 residues), poor representation of the reference sequence throughout the MSA, positions with many gaps, insufficient sequences in the MSA, or overly long sequence names. If you encounter problems accessing results from the GEMME/iGEMME servers or need to perform large-scale analysis on multiple proteins, we recommend downloading the Docker image, which includes all necessary dependencies. Most of these issues can be resolved; for example, the number of sequences in the MSA can be adjusted to exceed 40,000. Please note that processing MSAs with a large number of sequences or very long reference sequences may take more time.

### 3.4 Computational time

GEMME/iGEMME is very fast since it does not rely on any training due to learning required by the methodological approach. The computational evaluation for GEMME is found in [1] and [9] and for iGEMME in [10].

### 3.5 Additional inputs in subsequent developments

Users can enhance predictions by utilizing additional information available through ESCOTT and PRESCOTT [10]. ESCOTT takes as additional input a 3D experimental or predicted structure, and PRESCOTT takes allele frequency data [10]. ESCOTT and PRESCOTT follow the AVE Alliance guidelines for variant effect predictors [43].

### 3.6 Compliance with the Atlas of Variant Effects (AVE) Alliance guidelines

GEMME/iGEMME follows several guidelines from the AVE Alliance for variant effect predictors [43]. Specifically, GEMME/iGEMME codebase is publicly accessible, clearly documented, and open-source. GEMME/ iGEMME algorithm is transparent, making it possible to rationalise the predictions with respect to the input sequence data. Moreover, since GEMME/iGEMME is not based on learning, it avoids circularity issues. We made GEMME/iGEMME predictions Findable, Accessible, Interoperable, and Reusable (FAIR) by the community. They include human proteome-wide scores and also scores computed on ProteinGym [2] to facilitate independent benchmarking and analysis. GEMME/iGEMME scores can be easily scaled between 0 and 1 for facilitating comparison with other variant effect predictors. Finally, GEMME/iGEMME is population-free, *i*.*e*. it does not use any human population or clinical variants, thus avoiding pervasive ancestry bias [44].

### 3.7 Software availability

GEMME is available online at http://www.lcqb.upmc.fr/gemme/ under the MIT Free Software License as a stand alone package, a Docker image (https://hub.docker.com/r/elodielaine/gemme) and a webserver. All GEMME parameters can be modified depending on users needs through configuration files directly in the Docker image. The webserver default parameters are set so as to ensure rapid execution – the number of iterations and of sequences for conservation calculation can be modified in the submission form. In addition, we provide public access to GEMME predictions of single mutational landscapes for the entire human proteome (20 586 proteins from the Swiss-Prot reviewed human proteome available in UniProt, as of August 2023) on Dryad [32].

iGEMME is accessible online at http://prescott.lcqb.upmc.fr, by providing to the ESCOTT web-server a sequence alignment without structure. ESCOTT and PRESCOTT are available at http://prescott.lcqb.upmc.fr, together with a database of ESCOTT precomputed prediction for more than 19 000 human proteins, accessible for visualisation. iGEMME and ESCOTT are a part of the PRESCOTT software; a docker image of PRESCOTT is available at https://hub.docker.com/repository/docker/tekpinar/PRESCOTT-docker/general and the source code is provided at http://gitlab.lcqb.upmc.fr/tekpinar/PRESCOTT under the CC BY-NC-SA 4.0 International license.

